# Plasticity-Driven Self-Organization under Topological Constraints Accounts for Non-Random Features of Cortical Synaptic Wiring

**DOI:** 10.1101/027268

**Authors:** Daniel Miner, Jochen Triesch

**Author notes:** This author performed the programming, analysis, and writing, and contributed to the conceptualization (Daniel Miner). This author provided significant expertise and consultation, provided initial editing, and contributed to the conceptualization (Jochen Triesch).

## Abstract

Understanding the structure and dynamics of cortical connectivity is vital to understanding cortical function. Experimental data strongly suggest that local recurrent connectivity in the cortex is significantly non-random, exhibiting, for example, above-chance bidirectionality and an overrepresentation of certain triangular motifs. Additional evidence suggests a significant distance dependency to connectivity over a local scale of a few hundred microns, and particular patterns of synaptic turnover dynamics, including a heavy-tailed distribution of synaptic efficacies, a power law distribution of synaptic lifetimes, and a tendency for stronger synapses to be more stable over time. Understanding how many of these non-random features simultaneously arise would provide valuable insights into the development and function of the cortex. While previous work has modeled some of the individual features of local cortical wiring, there is no model that begins to comprehensively account for all of them. We present a spiking network model of a rodent Layer 5 cortical slice which, via the interactions of a few simple biologically motivated intrinsic, synaptic, and structural plasticity mechanisms, qualitatively reproduces these non-random effects when combined with simple topological constraints. Our model suggests that mechanisms of self-organization arising from a small number of plasticity rules provide a parsimonious explanation for numerous experimentally observed non-random features of recurrent cortical wiring. Interestingly, similar mechanisms have been shown to endow recurrent networks with powerful learning abilities, suggesting that these mechanism are central to understanding both structure and function of cortical synaptic wiring.

**Author Summary:** The problem of how the brain wires itself up has important implications for the understanding of both brain development and cognition. The microscopic structure of the circuits of the adult neocortex, often considered the seat of our highest cognitive abilities, is still poorly understood. Recent experiments have provided a first set of findings on the structural features of these circuits, but it is unknown how these features come about and how they are maintained. Here we present a neural network model that shows how these features might come about. It gives rise to numerous connectivity features, which have been observed in experiments, but never before simultaneously produced by a single model. Our model explains the development of these structural features as the result of a process of self-organization. The results imply that only a few simple mechanisms and constraints are required to produce, at least to the first approximation, various characteristic features of a typical fragment of brain microcircuitry. In the absence of any of these mechanisms, simultaneous production of all desired features fails, suggesting a minimal set of necessary mechanisms for their production.

## Introduction

The patterns of synaptic connectivity in our brains are thought to be the neurophysiological substrate of our memories, and framework upon which our cognitive functions are computed. Understanding the development of micro-structure in the cortex has significant implications for the understanding of both developmental and cognitive / computational processes. Such insight would be invaluable in understanding the root causes of cognitive and developmental impairments, as well as understanding better the nature of the computations realized by the cortex. It is believed that a small population of strong synapses forms a relatively stable backbone in recurrent cortical networks – perhaps the basis of long-term memories – while a larger population of weaker connections forms a more dynamic pool with a high rate of turnover [1–3]. It has been shown that much of the lateral recurrent connectivity of the layers of the cortex is significantly non-random [4–6], with a focus on layer 5 (L5), as this is more conventionally examined via slice studies. It is an open question which non-random features are developed as a result of direct genetic programming, neural plasticity under structured input, and spontaneous self-organization. We examine here several noted non-random features of recurrent cortical wiring that we believe can be explained as the result of spontaneous self-organization — specifically, self-organization driven by the interaction of multiple neural plasticity mechanisms. The features we will examine are the heavy-tailed, log-normal-like distribution of synaptic efficacies or dendritic spine sizes [6–10] and their associated synaptic dynamics, and the overrepresentation of bidirectional connectivity and certain triangular graph motifs [6].

The interaction of multiple plasticity mechanisms, such as synaptic scaling and Hebbian plasticity has been studied before [11–14], with results suggesting that the interactions for such mechanisms are useful for the formation and stability of patterns of representation. However, we desire a more detailed look at how such self-organization might take place in the cortex. The predecessor to the model we use to address these issues is the Self-Organizing Recurrent Neural Network, or SORN [11]. The SORN is a recurrent network model of excitatory and inhibitory binary neurons which incorporates both Hebbian and homeostatic plasticity mechanisms. Specifically, it incorporates binarized spike timing dependent plasticity (STDP), synaptic normalization (SN), and intrinsic homeostatic plasticity (IP). Certain variants also employ structural plasticity. It has been demonstrated to be computationally powerful and flexible for unsupervised sequence and pattern learning, presenting apparent approximate Bayesian inference and sampling-like behavior [15–17]. Additionally, it has been used to reproduce synaptic weight distributions and growth dynamics observed in the cortex [18].

In this paper, we introduce the LIF-SORN, a leaky integrate-and-fire based SORN-inspired network model that incorporates a spatial topology with a distance-dependent connection probability, in addition to more biologically plausible variants of and extensions to the plasticity mechanisms of the SORN. The LIF-SORN models a recurrently connected network of excitatory and inhibitory neurons in L5 of the neocortex, or a slice thereof. This new model is the first to reproduce numerous elements of the synaptic phenomena examined in [10], [19], and [18] in combination with the sort of non-random graph connectivity phenomena observed in [6]. The simultaneous reproduction of all these elements with a minimal set of plasticity mechanisms and constraints represents an unprecedented success in explaining noted features of the cortical micro-connectome in terms of self-organization.

## Materials and Methods

### Simulation Methods

We randomly populate a 1000 x 1000 *μ*m grid with 400 LIF neurons with intrinsic Ornstein-Uhlenbeck membrane noise as the excitatory pool, and a similar (though faster refracting) population of 80 noisy LIF neurons as the inhibitory pool. All synapses are inserted into the network with a gaussian distance-dependent connection probability profile with a half-width of 200 *μ*m. This particular profile is chosen as a middle ground between the results of [6], which finds no distance dependence up to a scale of 80 - 100 *μ*m, and the results of [5], which finds an exponential distance dependence at a scale of 200 - 300 *μ*m. Recurrent excitatory synapses are not populated, as they will be grown “naturally” with the structural plasticity. Excitatory to inhibitory and inhibitory to excitatory synapses are populated to a connection fraction of 0.1 and inhibitory recurrent synapses to a connection fraction of 0.5, in approximate accordance with L5 experimental data [20]. Excitatory to inhibitory, inhibitory to excitatory, and inhibitory to inhibitory connections are given fixed efficacies and connectivities. Recurrent excitatory connectivity is begun empty and is to be grown in the course of the simulation. The relevant parameters are summarized in Tables 1 and 2.

**Table 1.** Basic network parameters.

| parameter | value |
| --- | --- |
| $N_{\text{exc}}$ | 400 |
| $N_{\text{inh}}$ | 80 |
| sheet size | $1000 \times 1000 \mu\text{m}$ |
| connection probability profile | $200 \mu\text{m}$ half-width Gaussian |

**Table 2.** Basic connectivity parameters. ^*^ indicates growth via plasticity

| parameter | EE | EI | IE | II |
| --- | --- | --- | --- | --- |
| connection fraction | target 0.1* | 0.1 | 0.1 | 0.5 |
| connection strength at insertion | 0.0001 mV* | 1.5 mV | -1.5 mV | -1.5 mV |
| conduction delay | 1.5 ms | 0.5 ms | 1.0 ms | 1.0 ms |

We use the Brian spiking neural network simulator [21]. The neuron model is a leaky integrate-and-fire (LIF) neuron, the behavior of which is defined by:

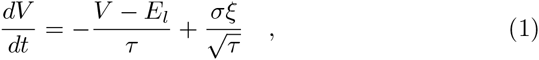

where *V* is the membrane potential, *E*_l_ is the resting membrane potential, *τ* is the membrane time constant, *σ* is the standard deviation of the intrinsic membrane noise, and *ξ* is the Ornstein-Uhlenbeck process which drives the noise. In our model, the variance of the noise is 5 mV. When V reaches a threshold *V*_T_, the neuron spikes, and the membrane potential *V* is returned to *V*_reset_ (which may be lower than *E*_l_ in order to provide effective refractoriness). The parameters used are given in Table 3.

**Table 3.** LIF neuron parameters.

| parameter | value |
| --- | --- |
| $E_l$ | -60 mV |
| $\tau$ | 20 ms |
| $V_{\text{reset}}^{\text{exc}}$ | -70 mV |
| $V_{\text{reset}}^{\text{inh}}$ | -60 mV |
| $\sigma$ | $\sqrt{5}$ mV |
| $V_T$ | variable via IP |

All parameters are shared between excitatory and inhibitory units unless otherwise denoted by superscripts “exc” and “inh.”

A simple transmitting synapse model is used, connecting neuron *i* to neuron *j*. When neuron *i* spikes, the synaptic weight *W_ij_* is added to the membrane potential *V* of neuron *j* following the conduction delay for the type of connection (as in Table 2). To improve network activity stability, this synaptic weight is modulated by a short term plasticity (STP) mechanism [22] implementing a rapid synaptic depression combined with a slightly slower facilitation. The STP mechanism consists of a two variable system:

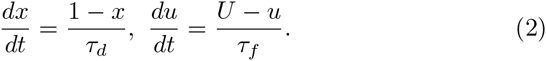

Upon each presynaptic spike, the variables are updated according to the following rules:

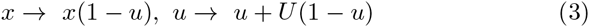

The synaptic weight is then modulated as 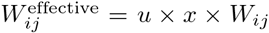. We select *U* = 0.04, *τ_d_* = 500 ms and *τ_f_* = 2000 ms as the respective depression and facilitation timescales, corresponding to approximate experimentally observed values [22, 23]. The presence of the STP adds a significant degree of stability to network activity and provides a more robust paramter range for other mechanisms, reducing the need for parameter tuning.

As in the original binary SORN, we include multiple plasticity mechanisms. The first is exponential spike timing dependent plasticity (STDP), which is executed at a biologically realistic timescale [24–29]. This defines the weight change to a synapse caused by a pair of pre- and post-synaptic spikes as in Equations 4, 5, and 6:

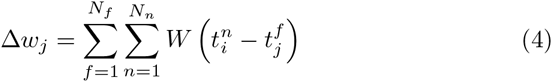

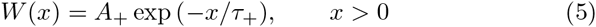

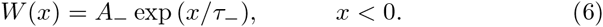

Here, *j* indexes the synapse, *f* indexes presynaptic spikes, and *n* indexes postsynaptic spikes. *A*_+_ and *A*_–_ are the maximal amplitudes of the weight changes, and *τ*_+_ and *τ*_–_ are the time constants of the decay windows. Values are set to approximate experimental data; in particular, round numbers were chosen that roughly approximate data in [24] and [25], with *τ*_+_ = 15 ms, *A*_+_ = 15 mV, *τ*_–_ = 30 ms, and *A*_–_ = 7.5 mV. We use the “nearest neighbor” approximation in order to efficiently implement this online, in which only the closest pairs of pre- and post-synaptic spikes are used. This is implemented in an event-based fashion, using a spike memory buffer with a timestep equal to that of the simulation itself (0.1 ms) and with the full calculation only evaluated upon a spike.

In the brain, several mechanisms appear to regulate the amount of synaptic drive that a neuron is receiving. [30] demonstrated the phenomenon of synaptic normalization during long-term potentiation (LTP). The summed areas of the synaptic active zones per micrometer of dendrite stay roughly constant, but the active zone area increases for some synapses while the total number of synapses per micrometer of dendrite decreases. This suggests that synaptic efficacies are mainly redistributed over the dendritic tree during the typical time course of an LTP experiment, but the sum of these efficacies (roughly corresponding to the sum of the active zone areas) stays approximately constant. Another phenomenon regulating the synaptic drive a neuron is receiving is homeostatic synaptic scaling [31], which is thought to regulate synaptic efficacies in a multiplicative fashion on a very slow time scale (on the order of days) in order to maintain a certain desired level of neural activity. For the sake of simplicity, we use here only a multiplicative form of normalization that drives the sum of synaptic efficacies to a desired target value on a fast time scale:

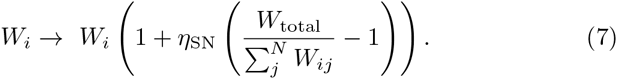

Here, *W*_i_ is the vector of incoming weights for any neuron *i*, *W_i_j* are the weights of the individual synapses, *W*_total_ is the target total input for each neuron, and *η*_SN_ is a rate variable which, together with the size of the timestep, determines the timescale of the normalization. *W*_total_ is calculated before the simulation run for each of the four types of synapse (E to E, E to I, I to E, and I to I) by multiplying the connection fraction for that type of connection by the mean synapse strength and the size of the incoming neuron population. The timescale we use is on the order of seconds and therefore accelerated from biology; corresponding to an application of the process once per second and *η*_SN_ = 1.0. We have tested it as well applying the normalization at every single simulation timestep, and with smaller values for *η*_SN_, which, except for very small values of *η*_SN_, has no significant effect on any of our observables. The accelerated timescale is sufficiently separated from that of the STDP, which operates on the order to tens of milliseconds, to avoid unwanted interactions while decreasing the necessary simulation time.

Neuronal excitability is regulated by various mechanisms and over different time scales in the brain. On a very fast milliseconds time scale, a neuron’s refractory mechanism prevents it from exhibiting excessive activity in response to very strong inputs [32]. This is inherently included in the neuron model we use. At a somewhat slower time scale, spike rate adaptation reduces many types of neurons’ responses to continuous drive [33]. Given that our model lacks any strong external drive, we neglect this. At very slow time scales of hours to days, intrinsic plasticity mechanisms change a neuron’s excitability through the modification of voltage gated ion channels that can modify its firing threshold and the slope of its frequency-current curve in a homeostatic fashion. Additional regulation of neuronal activity has been observed over multiple timescales [34, 35]. In order to capture the essence of such mechanisms in a simple fashion, we adopt a simple regulatory mechanism for the firing threshold, which, in combination with the previously discussed STP mechanism, phenomenologically captures the majority of these adaptive behaviors over short and medium timescales. Though relatively stable network activity can be achieved without this mechanism, it requires hand tuning of thresholds dependent on other network parameters, which we wish to avoid. The mechanism is implemented at discrete time steps in the following way:

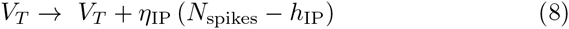

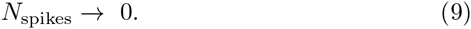

Here, *V_T_* is the threshold for an individual neuron, *η*_IP_ is a learning rate, *h*_IP_ is the target number of spikes per update interval, and *N*_spikes_ is the number of times a neuron has spiked since the last time a homeostatic plasticity step was executed. The right arrow indicates that the counter is reset after each evaluation of the window. This operation is performed at a biologically accelerated timescale. The desired target rate is chosen to be 3.0 Hz, so *h*_IP_ = 3.0 Hz x 0.1 ms = 0.0003 and *η*_IP_ is set to 0.1 mV. In our implementation, the operation is performed at every timestep of the simulation (0.1 ms), so *N*_spikes_ effectively becomes a binary variable and 9 becomes irrelevant. In this case, the action of the mechanism is that every spike increases the threshold by a small amount, and the absence of a spike decreases it by a small amount. Like the SN process, the accelerated (relative to biology) timescale is sufficiently separated from the timescale of the STDP to avoid unwanted interactions while decreasing the necessary simulation time.

We implement structural plasticity for the recurrent excitatory synapses via simultaneous synaptic pruning and synaptic growth. Synaptic pruning is implemented in a direct fashion in which synapses whose strengths has been driven below a near-zero threshold (0.000001 mV) by the other plasticity mechanisms are eliminated. At the same time, new synapses are stochastically added with a strength of 0.0001 mV, according to the distance-dependent per-pair connection probabilities, at a regular rate. This is done at an accelerated timescale by adding a random number of synapses (drawn from an appropriately scaled normal distribution) once a second. A mean growth rate is hand-tuned to lead to the desired excitatory-excitatory connection fraction. In this case, the mean growth rate is 920 synapses per second (with standard deviation of 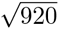) and the target connection fraction is 0.1 [6, 20]. The synapses are added according to pre-calculated connection probabilities determined by the gaussian connectivity profile described in the first paragraph of this section. Like the previous two plasticity mechanisms, the acceleration of the timescale from biology is justified by the principle of separation of timescales. At certain points in the Results and Supplementary Material, the results of the simulation are compared to those of a purely topological network. This is generated simply by performing the batch structural growth operation, as described, a single time, but adding instead a number of connections equal to the total number of connections at the target connection fraction.

## Results

### Network growth and abundance of bidirectional connections

As the fully simulated network runs, new recurrent excitatory synapses are allowed to grow and, if their strengths are driven close to zero, be pruned. The network first enters a growth phase, which lasts 100-200 seconds of simulation time, and then a stable phase in which the growth rate balances the pruning rate. The network is allowed to run for 500 seconds and the state of the excitatory connectivity and the dynamics of the connection changes during the final epoch are then examined.

We first examine, alongside the smooth growth of the network, the prevalence of bidirectional connections as compared to chance, a phenomenon noted to be significantly above-chance in [4] and [6], as shown in Figure 1. We observe for the total connection fraction a reliable value of 0.1, as selected. We observe a stable phase value of 0.018 for the bidirectional connection fraction, translating to a factor of 1.83 above chance. Our control for chance is the expected number of bidirectional connections for an Erdős-Rényi graph containing the same number of nodes and edges as the simulated network. For comparison purposes, a value of approximately 4 times chance is observed in [6]. We note that an otherwise equivalent non-topological network, in which the probability of connection between neurons is uniform rather than distance-dependent, produces a slight underrepresentation of bidirectional connections, reinforcing the well-known expectation that classical STDP, in the absence of other factors, favors unidirectional connectivity.

**Figure 1.**
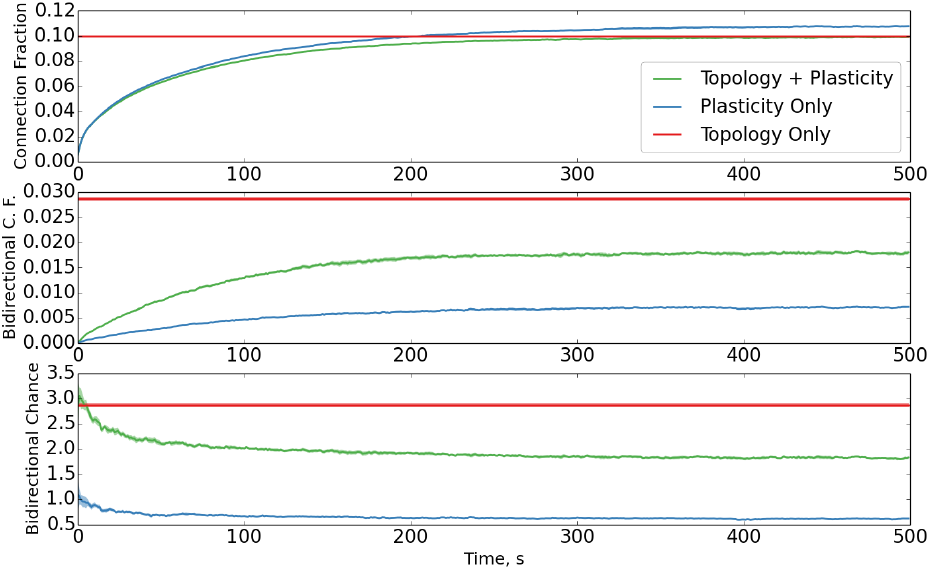
Evolution of total and bidirectional connection fraction with simulation time. Connection fraction evolution for plastic networks with and without topology, as well as flat values for topology only. (top) Growth and subsequent stabilization of the connection fraction of the network with simulation time. (middle) Growth of the bidirectional connection fraction. (bottom) Evolution of the bidirectional connection fraction with time as it relates to chance level (i.e. compared to the value for an Erdos-Rényi graph with the same number of nodes and edges). Data averaged over ten trials; standard deviation is shaded.

Regarding the growth of the network and the stabilization of its activity, we note one additional thing. In Figure 2, we observe that the distribution of interspike intervals (ISIs) and their coefficients of variation (CVs) follow the properties of an approximately Poisson-like spiking with an effective refractory period, as is observed in cortical circuits. That is to say, the distribution of ISIs follows an exponential decay with a distortion, induced by the refractory period, at the low end, and that the CVs of the ISIs tend to be close to one.

**Figure 2.**
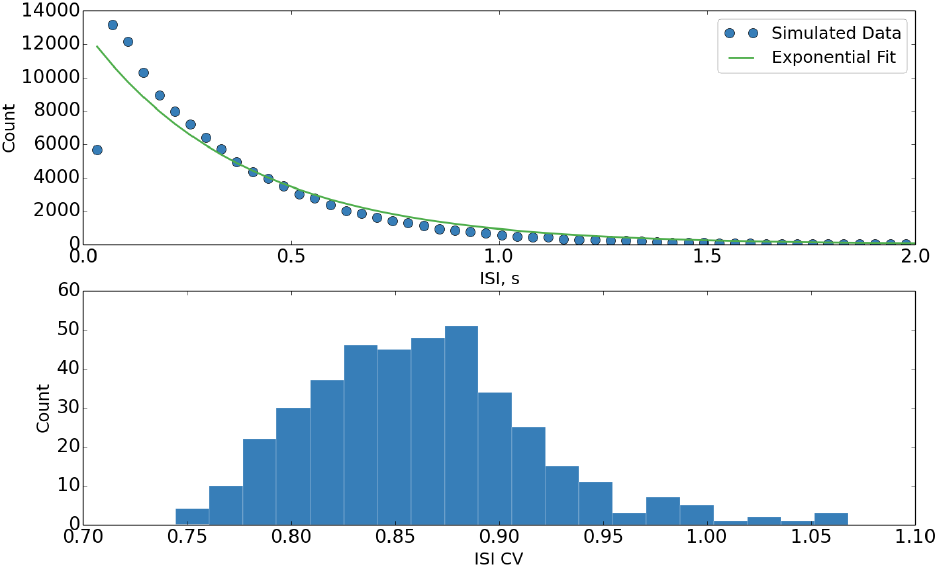
Distributions of ISIs and CVs thereof during stabilized network activity. (top) Pooled (over all neurons) distribution of ISIs with exponential fit, suggesting Poisson-like behavior with a refractory period. Individual neuron distributions have been tested to be similar. (bottom) Distribution of CVs of ISIs, suggesting Poisson-like behavior. Single trial data.

We would like to briefly consider how a model using classical STDP, which is known to drive the formation of unidirectional connections, can still produce such an abundance of bidirectional connections. In this model, the existence of clustering topology strongly drives the initial overrepresentation of bidirectional connections (as well as likely seeding higher order clustering effects, which are then selected and tuned via the plasticity mechanisms, and will be examined later). A simple mathematical argument will serve to demonstrate this (and, in fact, that any inhomogeneity in unidirectional connection probability will lead to an overrepresentation of bidirectional connections). Consider a single neuron in the center of a two dimensional sheet (this generalizes to volumes as well) which is populated with additional neurons at a uniform density. Assume that the central neuron has formed distance-dependent but otherwise random connections to the other neurons as follows: There is a local neighborhood containing a fraction *f* of all the neurons in the sampled area which have been connected with a high probability *p_h_*, while the remaining area contains the fraction 1 – *f* of all neurons, which connect with a lower probability *p_i_*. We can then treat the connection probability as a random variable *P* which takes the value *p_h_* with probability *f* and *p_l_* with probability 1 – *f* (this generalizes as well to additional neighborhoods, and, as the number of neighborhoods goes to infinity, to a continuous density of connection probability). The average overall connection probability of the neuron is then given by *E*[*P*] = *p_h_ f* + *p_l_* (1 – *f*). We now want to consider the average probability of finding a bidirectional connection. We assume that all neurons share the same distance-dependent connection probability, and therefore, the probability that a neuron within the local neighborhood has formed a connection to the central neuron is the same *p_h_* with which the central neuron is likely to form a connection to the neuron in the local neighborhood. Thus, the probability of a bidirectional connection in the local neighborhood is 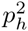, and by the same reasoning, the probability of forming a bidirectional connection with a neuron outside the local neighborhood is 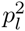. Then, the average overall bidirectional connection probability of the neuron is given by *E* [*P*^2^] = 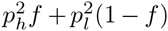. Since the squaring operation is convex, Jensen’s inequality applies, stating that for any convex function *g*(*P*) of a random variable *P*, *g* (*E*[*P*]) ≤ *E*[*g*(*P*)]. It then follows that with *g*(*P*) = *P*^2^, *E*[*P*^2^] ≥ *E*[*P*]^2^. Thus, bidirectional connections can occur more frequently than would be expected from the average unidirectional connection probability. Equality holds if and only if *P* is constant. It follows then that any inhomogeneity in unidirectional connection probability will lead to an overrepresentation of bidirectional connections. In the case of our model, the inhomogeneity is the distance-dependent connection probability, though any number of other factors could come into play.

For the above argument to apply to a structurally dynamic model such as ours, all that need be true is that bidirectional connections are added at a sufficiently high rate compared to their rate of removal due to STDP and pruning. The high number of bidirectional connections in the purely topological network, the low values for the purely plastic network, and the intermediate number of bidirectional connections for the full network model in Figure 1 serve to demonstrate the competition between the distance-dependent structural plasticity, which tends to boost bidirectional connectivity, and STDP and pruning, which tend to reduce bidirectional connectivity.

### Markov model of bidirectional overrepresentation

Furthermore, this competition can be captured and described by a simple Markov model in which each bidirectional connection pair develops independently of all the others.l. The model considers a pair of excitatory neurons and has three states {*U, S, D*} representing that the pair of neurons is either *unconnected, singly connected*, or *doubly connected*, respectively. We define transition probabilities denoting the probability of transitioning from one state to another during a fixed time interval. For example, *p_US_* is the probability for transitioning from the unconnected state *U* to the singly connected state *S*. The transition matrix is the matrix formed by all transition probabilites and is given by:

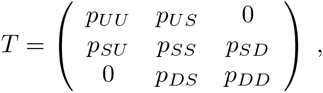

given the assumption that transitions from the unconnected state *U* to the doubly connected state *D* and vice versa are sufficiently unlikely to be considered negligible. Since the sum of the elements in each row of *T* has to equal one, *T* can be rewritten as:

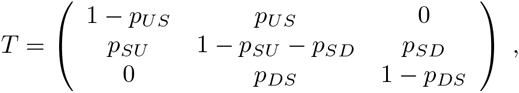

which depends on the four parameters *p_US_*, *p_SU_*, *p_SD_*, and *p_DS_*. If all of them are greater than zero, then the Markov Chain is regular and we can find its stationary distribution by finding the left Eigenvector of *T*:

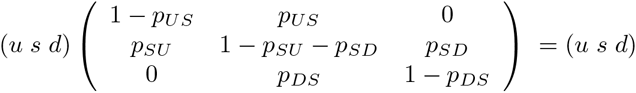

with *u* + *s* + *d* = 1. The resulting system of linear equations can be written as:

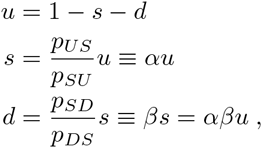

where we have defined *α* = *p_US_/p_SU_* and *β* = *p_SD_/p_DS_*. Thus, the behavior of the system depends only on the two transition probability ratios *α* and *β*. We can express *u* as a function of *α* and *β* to arrive at the final solution:

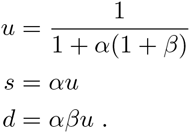

We can now consider the conditions under which the model leads to an overrepresentation of bidirectional connections. The overall connection probability in the Markov model is *p* = *s* / 2 + *d*. For a random graph, we then expect:

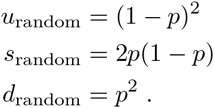

We consider an overrepresentation of bidirectional connections to be in comparison to a random graph. Therefore, using the previously defined transition ratios and a bit of algebra, we arrive the following expression for the overrepresentation *A*:

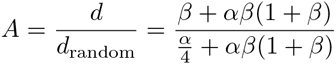

We can then empirically check this Markov model against our simulation. Counting and averaging connections and transitions over the last 100 seconds of a standard 500 second run of our model, we obtain *α* = 0.194 and *β* = 0.105. This leads the Markov model to predict an overrepresentation of *A* = 0.180, which is, in fact, the also measured value for the average overrepresentation over the observed time period.

### Statistics and fluctuations of synaptic efficacies

During the growth phase of the simulation, we note the reproduction of some of the results of [19], specifically that during network growth there is a tendency for larger synaptic weights to be more likely to shrink than smaller synaptic weights, as seen in Figure 3.

**Figure 3.**
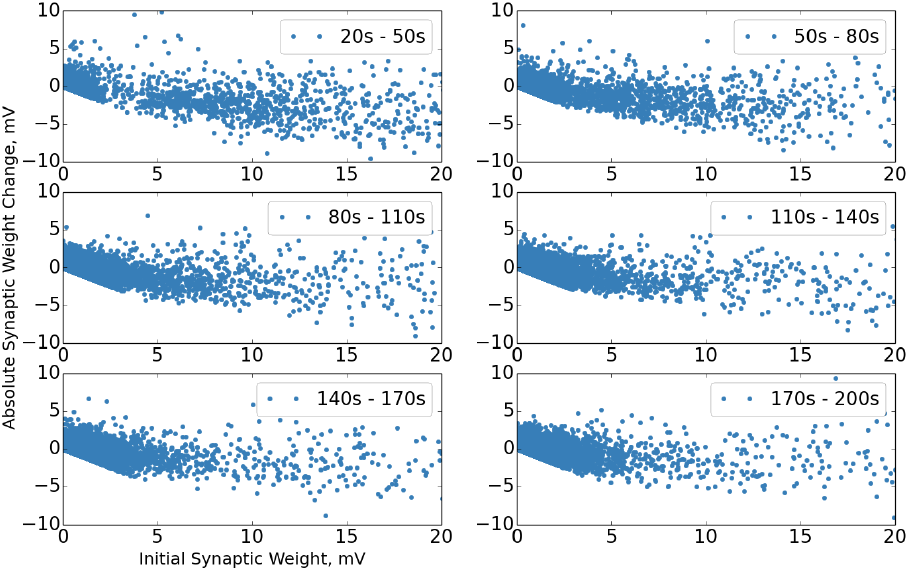
Synaptic change dynamics during network growth. Synaptic change dynamics during network growth epochs, before stabilization. Change is over entire epoch. “Bunching” in earliest epoch is an artifact of normalization under a small number of synapses. Single trial data.

Once the stable phase is reached, we observe the distribution of synaptic weights via histogramming, as previously stated, in Figure 4. This is in qualitative agreement with the heavy-tailed, log-normal-like shape typically observed in experimental data [6–10]. Several theoretical explanations for this distribution have been proposed, including a self-scaling rich-get-richer dynamic [18] and a confluence of additive and multiplicative processes [36, 37], both of which are consistent with our model. We note that the topology of the network seems to have a minimal effect on this result, as would be expected from the results of [18].

**Figure 4.**
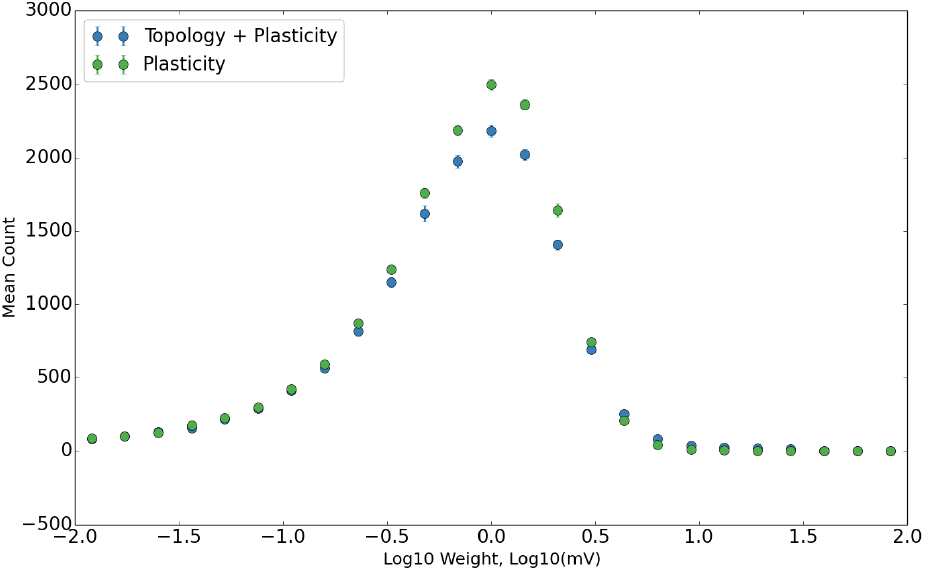
Log distribution of synaptic weights. The distribution of the base ten logarithm of synaptic weights for plastic networks with and without topology. Data averaged over ten trials; error bars are standard deviation.

We observe next the synaptic change dynamics in the stable phase of the network. We follow the format used in [10], comparing initial synaptic weight during a test epoch to both absolute and relative changes in synaptic weight, and demonstrate in Figure 5 that strong synaptic weights exhibit relatively smaller fluctuations over time, as experimentally observed [10]. Additionally, this serves to reinforce the earlier success of [18] in modeling such synaptic dynamics as the result of self-organization, and demonstrates that such results carry over into a biologically more realistic model.

**Figure 5.**
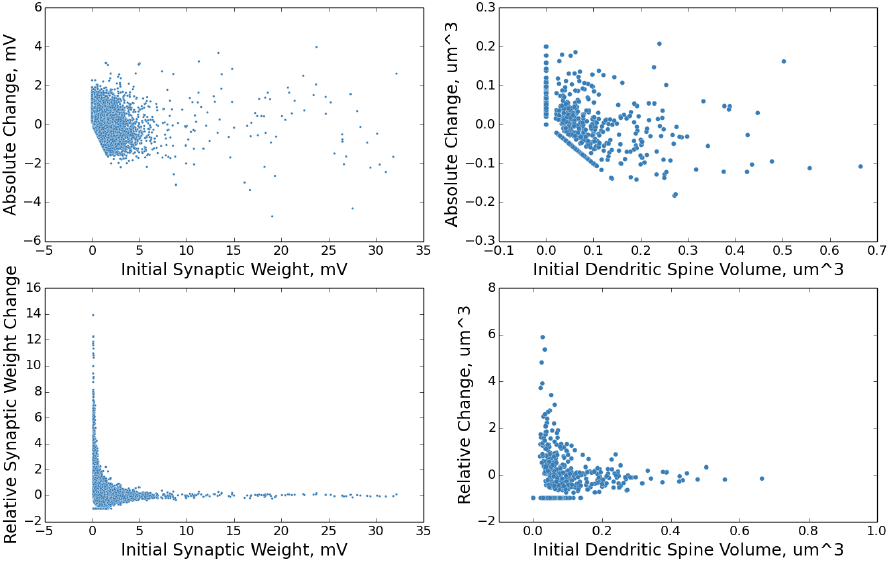
Change in synaptic weight as a function of initial synaptic weight. The above plots show the distributions of change in synaptic weight as a function of initial synaptic weight over a ten second simulation time period. The plots on the left are from the simulated network and are in electrophysiological units. The plots on the right are from experiment [10] and are in units of volume as estimated from fluorescence data. The plots on the top show the absolute change in synaptic weight / size. The plots on the bottom show the relative change in synaptic weight / size. Single trial data.

We examine, as well, the distribution of synaptic lifetimes. It has been predicted that the lifetimes of fluctuating synapses may follow a power law distribution [18]; our model makes this prediction as well. Recent experimental evidence supports this prediction [38]. We expand upon previous predictions with two interesting observations. In its current form, our model produces a slope of approximately 5/3 in the stable phase (for comparison, the experimentally observed slope is approximately 1.38). This decreases slightly in the growth phase. Secondly, we have observed as well that the slope can be modified by adjusting the balance of potentiation and depression in the STDP rule, varying between values between 1 and greater than 2, depending on the chosen parameters. For example, doubling the amplitude of the depression term in the STDP rule leads to a slope of approximately 5/2, while halving it leads to a slope of approximately 5/4. This is, in retrospect, an intuitive phenomenon. A preponderance of potentiation will lead to synapses being depressed to a value below the pruning threshold less frequently, thereby decreasing the slope of the power law. Similarly, in a depression-dominated scenario, synapses will be driven below the pruning threshold more frequently, leading to a higher power law slope. Returning to the slight decrease in slope during the growth phase, this makes sense, as a reduction in the effective pruning rate is necessary for the network to continue to grow. We believe that with a more extensive investigation of the effects of other model parameters on the power law, the slope of this distribution could be used as a meaningful measure of the potentiation-depression balance in a recurrent cortical network.

## Motif properties

We subsequently examine the prevalence of triadic motifs in the graph of the simulated network. An overrepresentation of certain motifs was noted in [6]. We used a script written for the NetworkX Python module [39, 40] to acquire a motif count for the graph of the simulated network. As the overrepresentation of bidirectional connections will trivially lead to an overrepresentation of graph motifs containing bidirectional edges, the control for chance is, in this case, a modified Erdős-Rényi graph with the same number of nodes, same number of unidirectional edges, and same number of bidirectional edges as the graph of the simulated network, with the unidirectional and bidirectional edges being independently populated. A similar control is used in [6]. We observe a similar pattern of “closed loop” triadic motifs being overrepresented in Figure 7, as experimentally observed in [6]. We note that the results for a non-topological plastic network with classical STDP, in the absence of additional factors, does not, relatively speaking, strongly select for any particular family of motifs. We similarly note that while distance-dependent topology does select for the observed family of motifs, it does not do so at the experimentally observed level. It is only the combination of topology and plasticity that strongly selected for the desired family of motifs while simultaneously producing all other noted effects. Approximate experimental data for comparison was extracted from [6] using GraphClick [41].

**Figure 7.**
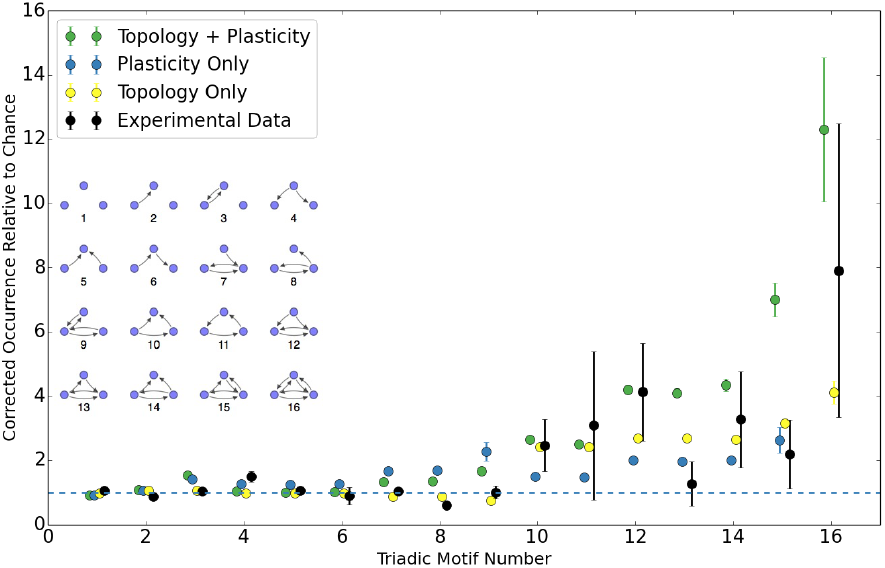
Triadic motif counts as a multiple of chance, corrected for bidirectional overrepresentation. Triadic motif counts (in the same order as [6]) for a simulated network as a multiple of chance value. The counts have been corrected for the observed overrepresentation of bidirectional connections. Results are shown for a complete network, a purely topological construction, an equivalent network with no topology, and approximate experimental data. For the topology-free network, the count of motif 16 is out of range due to the extremely low expected count after bidirectionality corrections. Data averaged over ten trials; error bars are standard deviation. Horizontal axis has been jittered slightly to increase readability.

## Discussion

The problem of how the non-random micro-connectivity of the cortex arises is a nontrivial one with significant implications for the understanding of both cognition and development. We attempt, in this paper, to provide insight into this problem by presenting a plausible model by which such non-random connectivity arises as the self-organized result of the interaction of multiple plasticity mechanisms under physiological constraints. Some models attempt to describe elements of the graph structure of the micro-connectome in purely physiological and topological terms [42]. However, such models necessarily lack an active network, and are thus unable to simultaneously account for synaptic dynamics, as our model does. Our model is, of course, a simple model, but the degree to which it accounts for observed non-random features of the typical cortical microcircuit without detailed structural features, metabolic factors, or structured input to drive the plasticity in a particular fashion is highly suggestive in terms of what is necessary at a bare minimum to drive the development and maintenance of the complex microstructure of the brain.

**Figure 6.**
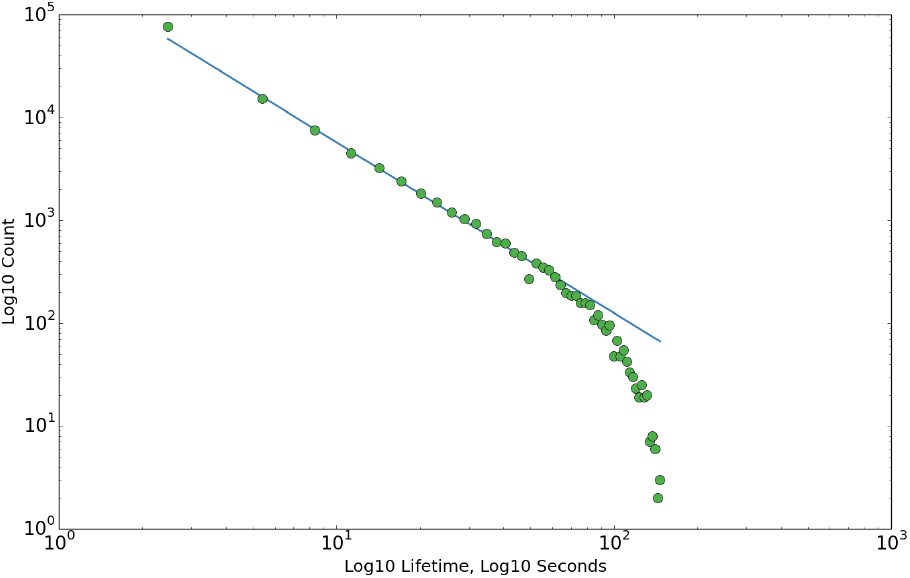
Distributions of synaptic lifetimes. The above plot shows the distributions of of synaptic lifetimes during the stable phase. Slope is approximately 5/3. The equivalent slope in the growth phase is slightly less. Here, we define entries in the growth phase as having synaptic end times of less than 150 seconds, and entries in the stable phase as having synaptic start times of greater than 350 seconds. Slopes are approximated via linear regression to the data points before the drop-off. Single trial data.

As mentioned in the introduction, it is hypothesized that a small backbone of strong synapses may form the stable backbone of long-term memory. The fact that in our model, strong weights remain stable in the presence of ongoing plasticity and despite significant fluctuations of smaller weights (which has been modeled as a stochastic Kesten process [37]), and the naturalness with which such a dynamic arises out of the interactions of known plasticity mechanisms, is both suggestive and supportive of this theory. On a related note, the heavy-tailed distribution of synaptic efficacies (often described as log-normal or log-normal-like) is an experimentally observed phenomenon seemingly fitting this narrative [6–10]. A theoretical explanation connecting log-normal firing rates with a log-normal synaptic efficacy distribution was one of the first proposed [43]. However, further studies have suggested that such a firing rate distribution is not necessary to create a heavy-tailed distribution of synaptic efficacies, using either a self-scaling rich-get-richer dynamic [18] or a combination of additive and multiplicative dynamics [36, 37].

An additional noted non-random feature of cortical recordings that has been passed over in this model is the observed log-normal distribution of cortical firing rates (touched upon in the previous paragraph). Our intrinsic plasticity mechanism necessarily negates this feature, which may be selforganized via mechanisms not included in our model, such as diffusive homeostasis [44, 45]. In order to maximize simplicity, a single target firing rate is chosen for all neurons. This also permits pooling of the ISIs for analysis. Additional testing in which the target firing rate is drawn from a log-normal distribution produces minimal qualitative effects on the observed features (except, trivially, the ISI distribution, see Figure S1). Another issue is that as things stand, the exact statistics of the micro-connectome are difficult to discern – though strong inferences can be made in the right direction – due to inherent sampling biases in paired patch-clamp reconstructions of limited size [46]. It is our hope and belief that advances in fluorescence imaging, automated electron microscopy reconstruction [47, 48], and massive multi-unit array recordings will help to alleviate these biases. One might imagine that additional biases may be caused by the relatively small model size of 400 excitatory neurons, when realistic cortical densities would result in thousands of neurons in such an equivalent volume. We have tested the network at much larger sizes of up to 2000 neurons and found no notable qualitative change to our observed results (Figure S2; all other features remain the same as well), so we maintained a relatively small network size to increase computational ease. It should be noted that except for this check, all supplementary checks, tests, and additional analyses were performed with the standard 400 + 80 neuron network size.

We have described the formation of the overrepresentation of bidirectional connections in terms of the competition between structural growth and structural pruning in the presence of a topological inhomogeneity. Other possibilities for increasing the prevalence of bidirectional connections include an STDP window with an integral greater than zero (i.e. biased toward potentiation), or one in which the asymmetries are finely tuned so that, given the target homeostatic target firing rate, connections are, on the whole, more likely to potentiate (making the STDP window fully symmetrical has, in our model, only a minimal effect). Additionally, more complicated STDP models [50, 51] are known to produce overrepresentation of bidirectional connections in high-frequency firing regimes.

One other computational study has reproduced similar motif overrepresentations, however, this model was significantly more complex and required specific structured input [49]. Some might view the fact that, in this model, the primary driver behind the overrepresentation of bidirectional connections is topology, as a shortcoming. We do not view this as a problem; after all, topology exists in the cortex and the rest of the study’s results suggest it is an important factor in the self-organization of cortical circuits. There are the previously mentioned mechanisms utilizing non-classical STDP, such as the so-called triplet and voltage rules [50, 51], which, in the presence of high-frequency activity, are capable of producing and maintaining bidirectional connections. Introducing such mechanisms into a similar model would be a welcome and interesting future study, and could potentially lead to an even stronger and more precise motif selectivity. To further explain the importance of the various mechanisms we have introduced in self-organization, we have included a brief analysis of the behavior of the network in the absence of individual mechanisms (see Table 4 below and Figures S3 and S4). Essentially, removal of the topology leaves the synaptic dynamics mostly unchanged, but significantly alters the connectivity structure. Removal of either element of structural plasticity, or of STDP (because without depression, no pruning will occur) lead to failure to form (in the case of growth) either divergent network growth (in the case of pruning or STDP). Removal of the STP leads to “epileptic” behavior, resulting in dynamic and structural disruptions. Removal of SN leads to a small subset of synapses experiencing runaway growth, with the others shrinking to near zero and being pruned. Finally, removal of the IP leads to small changes to the structural properties, but requires fine tuning of the thresholds to run even in this regime. Failure to tune the thresholds in this case leads to silent or epileptic networks.

**Table 4.**
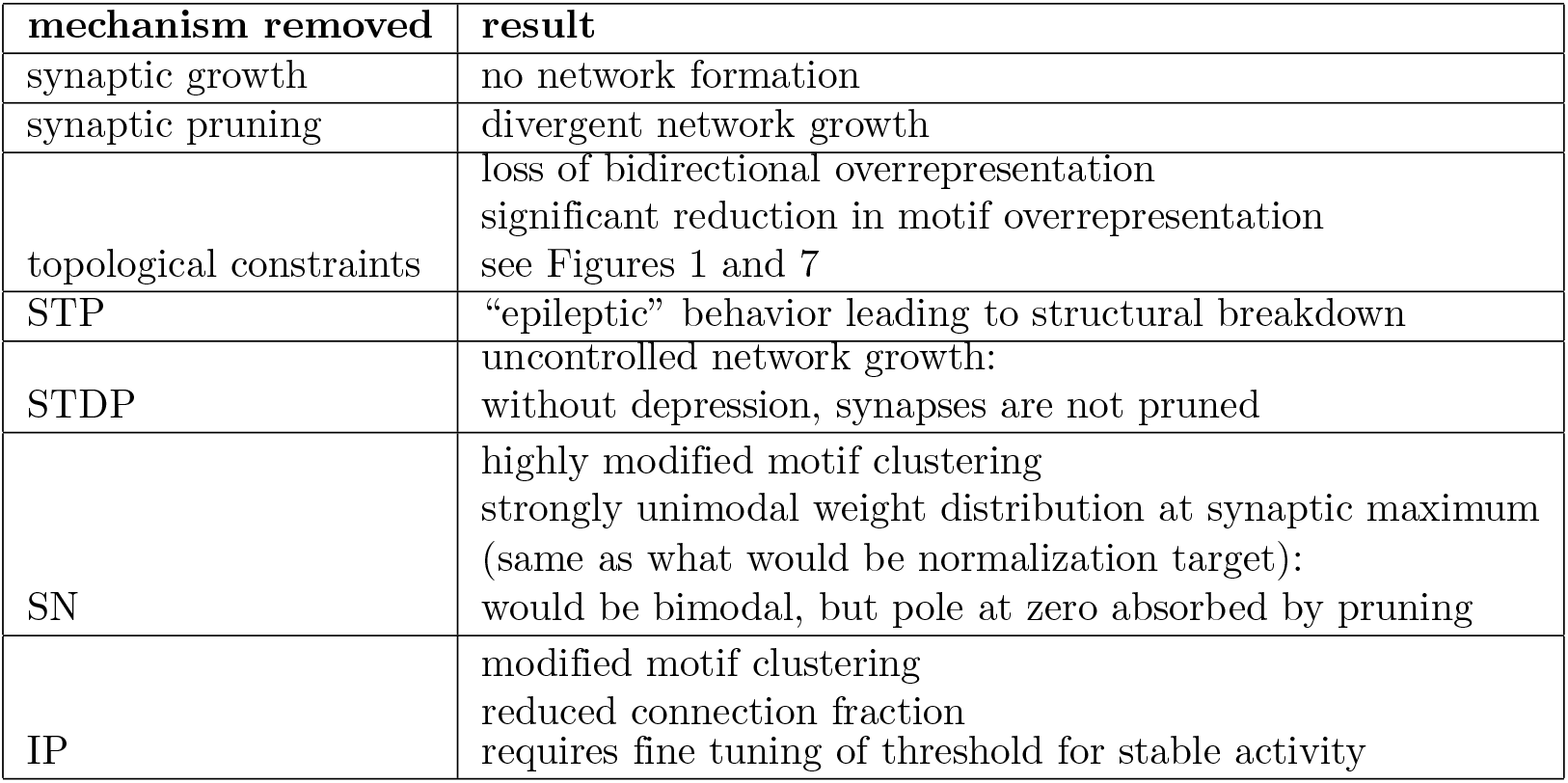
Results of plasticity mechanism removal. See Figures S4 and S3 for additional illustration.

| mechanism removed | result |
| --- | --- |
| synaptic growth | no network formation |
| synaptic pruning | divergent network growth |
| topological constraints | loss of bidirectional overrepresentation<br>significant reduction in motif overrepresentation<br>see Figures 1 and 7 |
| STP | “epileptic” behavior leading to structural breakdown |
| STDP | uncontrolled network growth:<br>without depression, synapses are not pruned |
| SN | highly modified motif clustering<br>strongly unimodal weight distribution at synaptic maximum<br>(same as what would be normalization target):<br>would be bimodal, but pole at zero absorbed by pruning |
| IP | modified motif clustering<br>reduced connection fraction<br>requires fine tuning of threshold for stable activity |

Additionally, with the aim of understanding the relationship between the activity correlation, the synaptic weights, and the intersomatic separation, a Spearman’s rank correlation analysis was performed on such data from an example trial (results in Table S1). In summary, a strong and highly significant positive correlation was found between the spike correlation and the synaptic weight, as would be expected from STDP. However, only a weak (negative) correlation was found between the spike correlation and the intersomatic separation, and no significant correlation was found between the intersomatic separation and synaptic weight.

As a concluding point, often, models of cortical microcircuits are described as random graphs, such as the classical random balanced network [52]. However, experiments have demonstrated that the structure of cortical microcircuitry is significantly non-random [5, 6], suggesting that random networks may be insufficient for modeling cortical development and activity. Lacking in structural plasticity of topology, such random graph based balanced networks are incapable of producing the sort of results we have observed. Having provided a mechanism with which one may generate a cortex-like non-random structure, it would be enlightening to determine if said structure provides any significant computational or metabolic advantage as compared to a random graph. Similarly, limitations in online plasticity capabilities significantly hinder the use of such random networks and their relatives in reservoir computing [53] for unsupervised learning and inference tasks (though progress has recently been made in this direction [54]), while earlier studies with the original SORN model [11, 15] suggest that the particular combination of plasticity mechanisms in our model can endow networks with impressive learning and inference capabilities. A logical next step is therefore to study the learning and inference capabilities of LIF-SORN networks and relate them to neurophysiological experiments. Our rapidly developing ability to manipulate neural circuits in vivo suggests this as an exciting direction for future research. It is our belief that the future of neural network-based computation and modeling of biological processes lies in the incorporation of multiple plasticity and homeostatic mechanisms under simple sets of constraints.

## Acknowledgments

We would like to thank the authors of [10] for sharing their data. We would like as well to thank Christoph Hartmann and Pengsheng Zheng for their valuable consultation.

## Supporting Information

**Supporting Tables**

**Table S1.** Spearman’s rank correlation and associated P-value between intersomatic separation, synaptic weight, and pairwise spike correlation. Representative single trial example data. Spike correlation was taken from 50 s activity with 50 ms bins [55].

| features | Spearman’s $\rho$ | P-value |
| --- | --- | --- |
| spike correlation and synaptic weight | $\rho = 0.32$ | $P = 0.00$ |
| spike correlation and separation | $\rho = -0.02$ | $P = 0.04$ |
| synaptic weight and separation | $\rho = -0.01$ | $P = 0.46$ |

**Supporting Figures**

**Figure S1.**
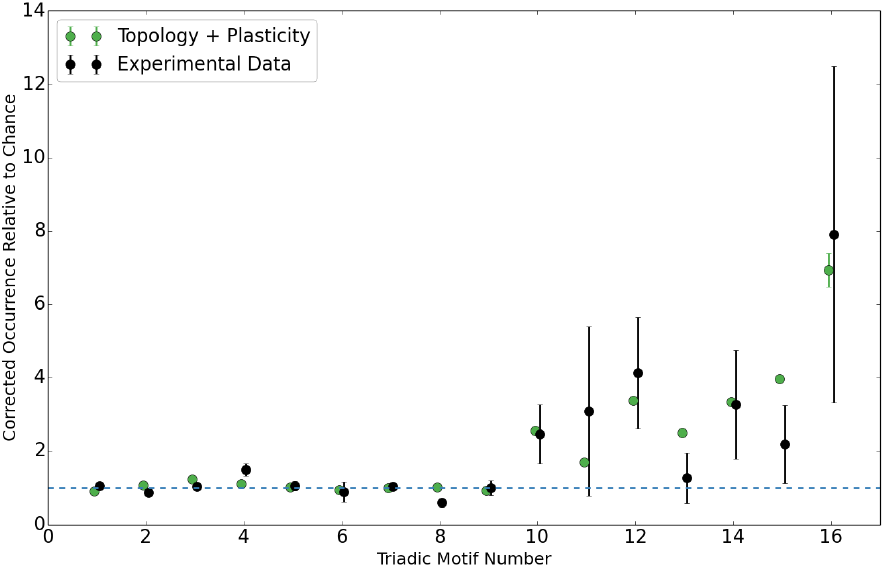
Triadic motif counts as a multiple of chance for lognormal firing rates, corrected for bidirectional overrepresentation. Triadic motif counts (in the same order as [6]) for a simulated network as a multiple of chance value. The counts have been corrected for the observed overrepresentation of bidirectional connections. Results are shown for a complete network with IP target rates drawn from a log-normal distribution (mean of 3.0, standard deviation of 1.0 Hz) instead of a single value and approximate experimental data.. Other parameters remain the same, aside from scaling of growth rate to obtain stable phase connection fraction of 0.1. Error bars are standard deviation. Horizontal axis has been jittered slightly to increase readability.

**Figure S2.**
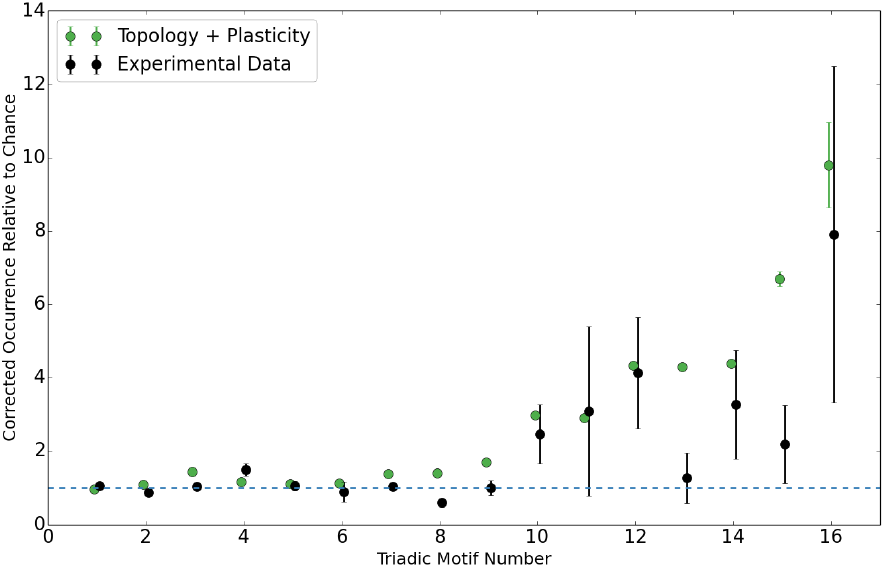
Triadic motif counts as a multiple of chance for a larger (2000 neuron) network, corrected for bidirectional overrepresentation. Triadic motif counts (in the same order as [6]) for a simulated network as a multiple of chance value. The counts have been corrected for the observed overrepresentation of bidirectional connections. Results are shown for a complete network of 2000 neurons and approximate experimental data.. Other parameters remain the same, aside from scaling of growth rate to obtain stable phase connection fraction of 0.1. Error bars are standard deviation. Horizontal axis has been jittered slightly to increase readability.

**Figure S3.**
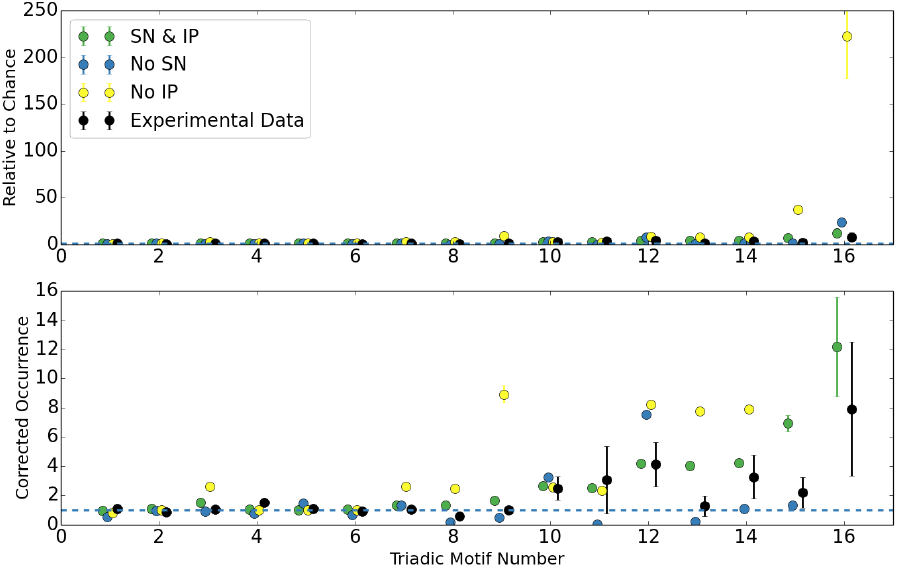
Triadic motif counts as a multiple of chance for networks with plasticity mechanisms removed, corrected for bidirectional overrepresentation. Triadic motif counts (in the same order as [6]) for a simulated network as a multiple of chance value. The counts have been corrected for the observed overrepresentation of bidirectional connections. Results are shown for a complete network, a network without IP, a network without SN, and approximate experimental data.. Error bars are standard deviation. Horizontal axis has been jittered slightly to increase readability.

**Figure S4.**
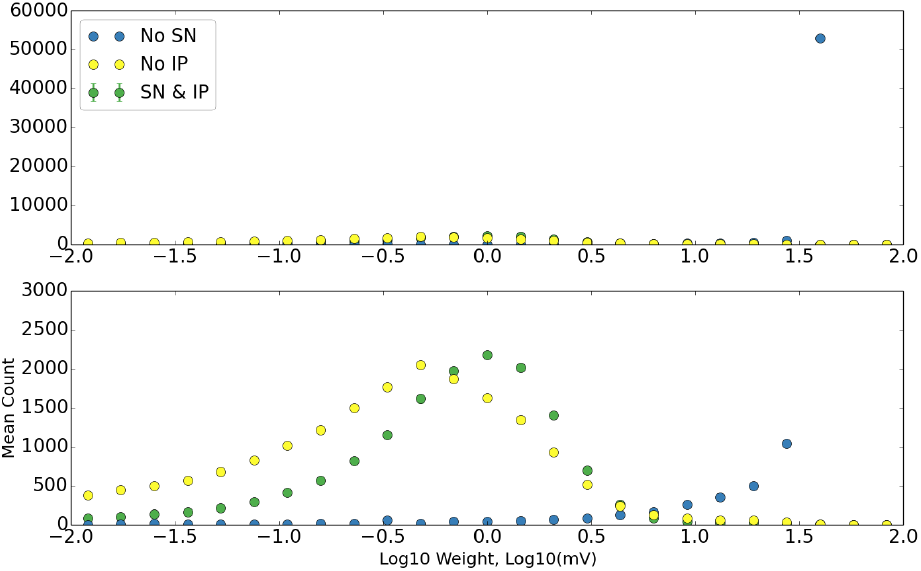
Log distribution of synaptic weights for networks with plasticity mechanisms removed. The distribution of the base ten logarithm of synaptic weights for a complete network (ten trials), a single network without IP, and a single network without SN. Error bars are standard deviation.

